# Behaviors and Energy Source of *Mycoplasma gallisepticum* Gliding

**DOI:** 10.1101/620922

**Authors:** Masaki Mizutani, Makoto Miyata

## Abstract

*Mycoplasma gallisepticum*, an avian-pathogenic bacterium, glides on host tissue surfaces by using a common motility system with *Mycoplasma pneumoniae*. In the present study, we observed and analyzed the gliding behaviors of *M. gallisepticum* in detail by using optical microscopes. *M. gallisepticum* glided at a speed of 0.27 ± 0.09 µm/s with directional changes relative to the cell axis of 0.6 ± 44.6 degrees/5 s without the rolling of the cell body. To examine the effects of viscosity on gliding, we analyzed the gliding behaviors under viscous environments. The gliding speed was constant in various concentrations of methylcellulose but was affected by Ficoll. To investigate the relationship between binding and gliding, we analyzed the inhibitory effects of sialyllactose on binding and gliding. The binding and gliding speed sigmoidally decreased with sialyllactose concentration, indicating the cooperative binding of the cell. To determine the direct energy source of gliding, we used a membrane-permeabilized ghost model. We permeabilized *M. gallisepticum* cells with Triton X-100 or Triton X-100 containing ATP and analyzed the gliding of permeabilized cells. The cells permeabilized with Triton X-100 did not show gliding; in contrast, the cells permeabilized with Triton X-100 containing ATP showed gliding at a speed of 0.014 ± 0.007 μm/s. These results indicate that the direct energy source for the gliding motility of *M. gallisepticum* is ATP.

**IMPORTANCE:** *Mycoplasmas*, the smallest bacteria, are parasitic and occasionally commensal. *Mycoplasma gallisepticum* is related to human pathogenic *Mycoplasmas*—*Mycoplasma pneumoniae* and *Mycoplasma genitalium*—which causes so-called ‘walking pneumonia’ and non-gonococcal urethritis, respectively. These *Mycoplasmas* trap sialylated oligosaccharides, which are common targets among influenza viruses, on host trachea or urinary tract surfaces and glide to enlarge the infected areas. Interestingly, this gliding motility is not related to other bacterial motilities or eukaryotic motilities. Here, we quantitatively analyze cell behaviors in gliding and clarify the direct energy source. The results provide clues for elucidating this unique motility mechanism.

## INTRODUCTION

Members of the bacterial class Mollicutes, including the genus *Mycoplasma*, are parasitic, occasionally commensal, and characterized by small cells and genomes as well as the absence of a peptidoglycan layer (1, 2). More than ten *Mycoplasma* species, such as the fish pathogen *Mycoplasma mobile* (3-5) and the human pathogen *Mycoplasma pneumoniae* (6-8), have membrane protrusions and exhibit gliding motility in the direction of the membrane protrusion on solid surfaces, which enables *Mycoplasmas* to parasitize higher animals.

Interestingly, *Mycoplasma* gliding does not involve flagella or pili and is entirely unrelated to other bacterial motility systems and the conventional motor proteins that are common in eukaryotic motility systems (9, 10).

The gliding motilities of *Mycoplasmas* are classified into two systems; *M. mobile*-type and *M. pneumoniae*-type (5, 8). *M. pneumoniae*-type gliding has until now been studied mainly in *M. pneumoniae* and *Mycoplasma genitalium*. A structure outline of the gliding machinery has been suggested, including that for fifteen component proteins (6-8, 11). The gliding machinery, called the ‘attachment organelle,’ is composed of an internal core and adhesin complexes (12-14). The internal core is divided into three parts, the bowl complex, paired plates, and terminal button (3, 6, 13-15). It has been suggested that the bowl complex connects the internal core to the cell body and may be responsible for the generation or transmission of force (8, 16, 17). Furthermore, paired plates are the scaffold for formation and force transmission of the gliding machinery (8, 18-24). The terminal button is thought to tightly bind to the front side of the cell membrane (8, 23, 25-27). The adhesin complex is composed of P1 adhesin and P90 (28). P90 is encoded in tandem with P1 adhesin and is cleaved from another protein, P40, for maturation (29, 30). A recent study shows that the homolog of P40/P90 in *M. genitalium*, P110, has a binding pocket of sialylated oligosaccharides (SOs) (31), which are binding targets for *Mycoplasma* infection and gliding. The mechanism of the gliding motility has been proposed as an ‘inchworm model,’ in which a cell catches SOs on solid surfaces through the adhesin complexes and is propelled by the repetitive extensions and retractions of the internal core (3, 7, 8).

The gliding motility of *M. mobile* is driven by ATP using ‘ghost’ which has a permeabilized membrane and can be reactivated by the addition of ATP (32-34). In contrast, the direct energy source for *M. pneumoniae*-type gliding motility is still unclear (8).

*Mycoplasma gallisepticum* is an avian pathogen that causes chronic respiratory disease in chickens and infectious sinusitis in turkeys. The cells transmit from breeder birds to their progeny *in ovo* (1, 35, 36). *M. gallisepticum* glides using the *M. pneumoniae*-type motility system and has eight homologs of component proteins of gliding machinery in *M. pneumoniae* whose identities for amino acids range from 20% to 45% (36-39). The structure of the gliding machinery is similar to that in *M. pneumoniae* (37, 39). *M. gallisepticum* has a faster growth rate and more stable cell shape than *M. pneumoniae*, which is beneficial for the motility study (37).

In this study, we observe and analyze the gliding behaviors of *M. gallisepticum* in detail, and clarify the direct energy source of the gliding motility by modified gliding ghost experiments.

## RESULTS

### Gliding behaviors

The gliding motility of *M. gallisepticum* has been reported previously (37, 40), but the details have not yet been examined. Therefore, these details were examined in this study. *M. gallisepticum* cells were collected, suspended in phosphate-buffered saline (PBS) containing 10% non-heat-inactivated horse serum, and inserted into a tunnel chamber constructed by taping coverslips and precoated with non-heat-inactivated horse serum and bovine serum albumin. Then, the tunnel chamber was washed with PBS containing 20 mM glucose (PBS/G) and was observed by phase-contrast microscopy. The cells showed a flask shape (Fig. 1A) and glided in the direction of the tapered end, as previously reported (Fig. 1B; see Movie S1 in the supplemental material) (37). The proportions of gliding cells to all cells and the gliding speeds averaged for 60 s at 1 s intervals were 62% ± 6% (n = 454) and 0.27 ± 0.09 μm/s (n = 231), respectively (Fig. 1B and C), which is consistent with those reported in the previous study (37). To analyze the gliding direction, we traced the angles between the cell axis and the following gliding direction for 60 s at 5 s intervals, as previously described (41). The averaged gliding direction relative to the cell axis was 0.6 ± 44.6 degrees/5 s (n = 231) (Fig. 1D), indicating that *M. gallisepticum* has no significant directional bias.

**Fig. 1.**
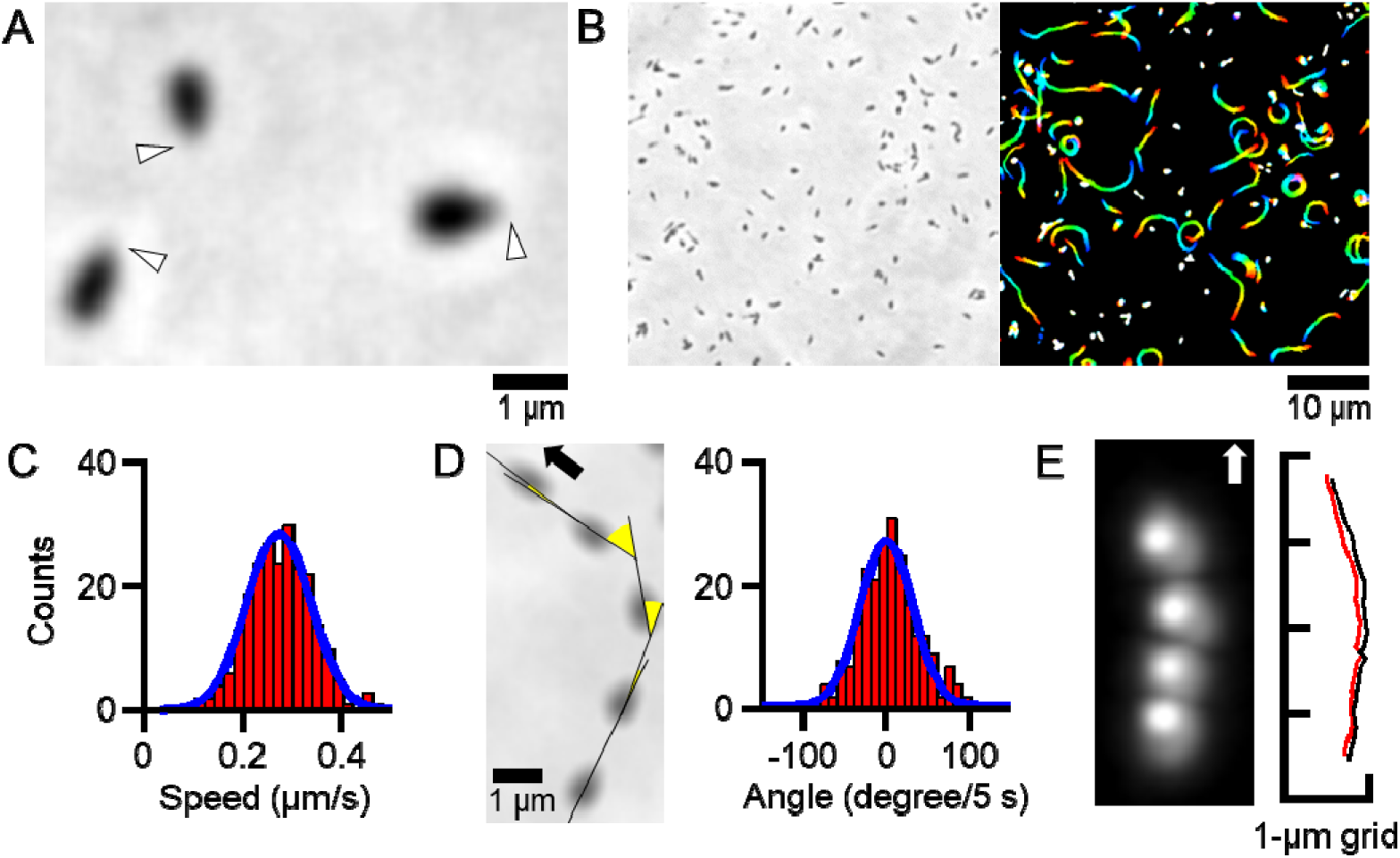
Gliding behaviors. (A) Phase-contrast micrograph of *M. gallisepticum* cells. The cells on the glass are gliding in the direction of the membrane protrusion marked by white triangles. (B) Field image of phase-contrast micrograph (left) and cell trajectories for 30 s, changing color with time from red to blue (right). (C) Distribution of gliding speeds averaged for 60 s at 1 s intervals was fitted with a Gaussian curve (n = 231). (D) Schematic illustration showing the measurement of gliding direction (left) and the distribution of gliding directions (right). Five consecutive cell images at 5 s intervals are shown in the same field. The cell axis and gliding directions are shown by black lines and yellow sectors, respectively. The cell glided in the direction indicated by the black arrow (left). The distribution was fitted with a Gaussian curve (right). (E) Dark-field micrograph of a cell attached with colloidal gold (left) and trajectories (right). Four consecutive images of a cell attached with colloidal gold at 10 s intervals are shown in the same field. The cell glided in the direction indicated by the white arrow (left). The trajectories of the mass centers of the cell and colloidal gold are indicated by black and red lines, respectively (right).

### Possibility of rolling around the cell axis

A previous study shows that the adhesin complexes of *M. pneumoniae* exist around the membrane protrusion (7). *M. gallisepticum* may have a similar distribution because it uses similar gliding machinery to *M. pneumoniae* (36-39). The distribution of adhesin complexes suggests that cells may roll around the cell axis during gliding. To examine this possibility, we traced the movement of 40 nm colloidal gold attached to a gliding cell. *M. gallisepticum* cells were biotinylated on the cell surface through amino groups, then mixed with 40 nm colloidal gold conjugated with streptavidin in the tunnel chamber and observed by dark-field microscopy. The cells attached with colloidal gold glided at a similar speed to cells without colloidal gold (see Fig. S1 and Movie S2 in the supplemental material). All pairs of mass centers of cells and colloidal gold moved while maintaining a constant distance (n = 20) (Fig. 1E; see Fig. S1 in the supplemental material), showing that the cells glide without rolling of the cell body.

### Gliding in viscous environments

A previous study shows that *M. mobile* gliding is drastically inhibited by viscous environments created using methylcellulose (MC) or Ficoll (41). To examine the effects of viscosity on *M. gallisepticum* gliding, we analyzed the gliding behaviors under viscous buffers including MC or Ficoll. MC is a long, linear, and slightly branched polymer, and forms a gel-like three-dimensional network. Ficoll is a highly branched polymer which increases viscosity and does not form a network (42, 43). *M. gallisepticum* cells were suspended in PBS/G containing various concentrations of MC or Ficoll, inserted into the tunnel chamber, observed by phase-contrast microscopy, and analyzed for gliding speed and direction (Fig. 2A). The gliding speed did not significantly change with an increase in viscosity from 0.22 ± 0.06 μm/s (n = 50) at 0.66 mPa s to 0.19 ± 0.04 μm/s (n = 50) at 7.3 mPa s with 0% and 0.50% MC, respectively. However, the gliding speed did significantly decrease to 0.11 ± 0.04 μm/s (n = 50) at 4.6 mPa s with 15% Ficoll (P<0.001 by Student’s *t*-test) (Fig. 2B and C). The averaged gliding direction relative to the cell axis did not change significantly with an increase in viscosity for all examined conditions. However, the standard deviations of gliding direction significantly decreased in Ficoll conditions (P<0.001 by F-test) from 27.4 degrees/5 s (n = 48) in 0% to 12.4 degrees/5 s (n = 50) in 15% Ficoll (Fig. 2B and C). These results indicate that the gliding motility of *M. gallisepticum* is affected by Ficoll but not MC.

**Fig. 2.**
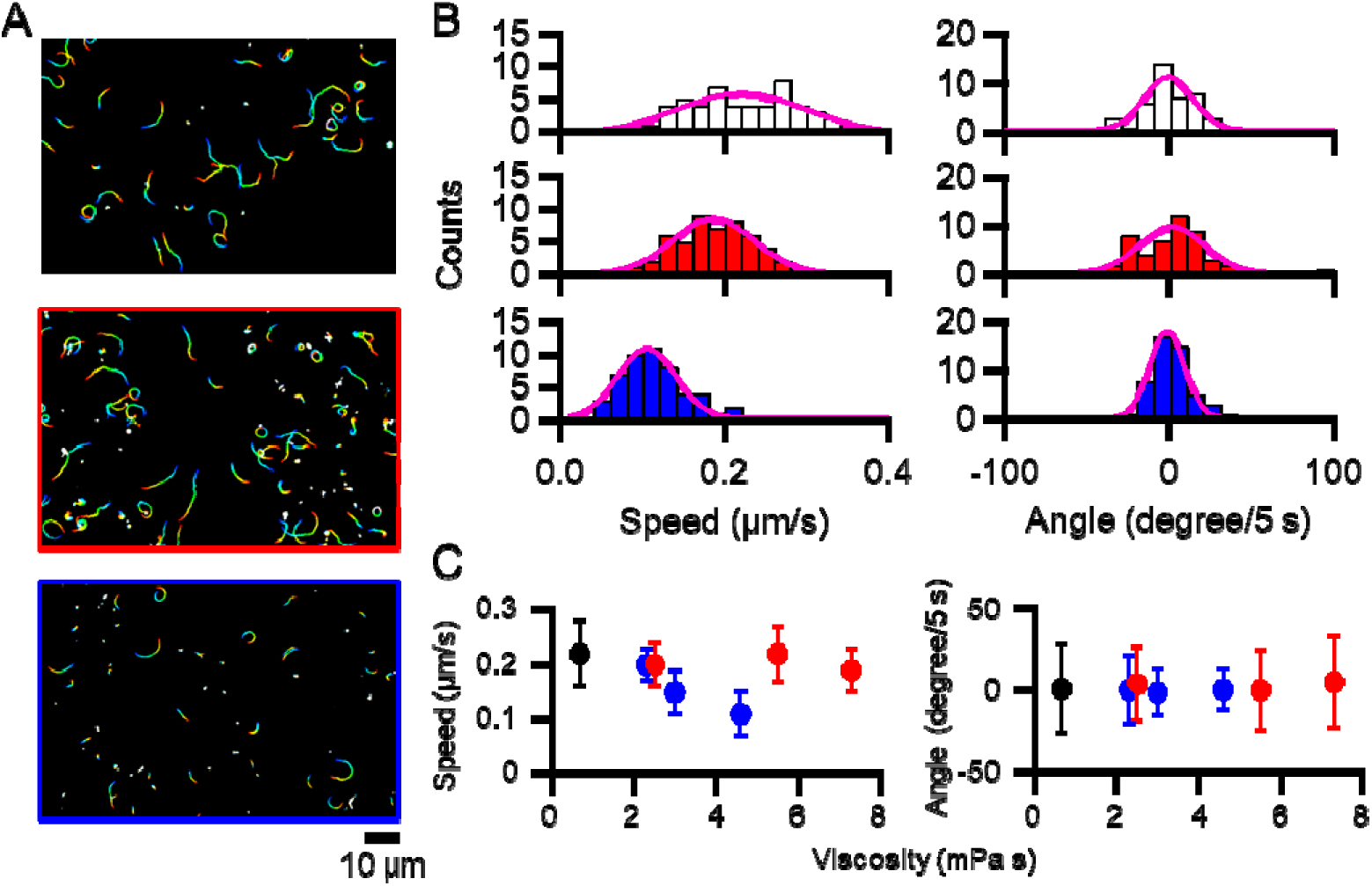
Effects of MC and Ficoll on gliding. (A) Cell trajectories for 60 s under PBS/G (top), 0.50% MC (middle) and 15% Ficoll (bottom). The color changes with time from red to blue. (B) Distributions of gliding speeds and directions under PBS/G (top), PBS/G containing 0.5% MC (middle) and 15% Ficoll (bottom) were fitted with Gaussian curves. (C) Gliding speeds and directions under various concentrations of MC or Ficoll. The average gliding speeds for 60 s at 1 s intervals and the average gliding directions every 5 s for 60 s under PBS/G containing 0%, 0.10%, 0.25%, 0.50% MC or 5%, 10%, 15% Ficoll conditions are plotted with standard deviations. Black, red, and blue circles indicate PBS/G, MC, and Ficoll, respectively.

### Inhibition of binding and gliding by free sialyllactose

Previous studies show that *M. mobile* glides via dozens of working legs, the numbers of which can be reduced by the addition of free SOs (33, 44-46). To examine the relationship between binding and gliding, we added various concentrations of free 3′-*N*-acetylneuraminyllactose (3′-sialyllactose, SL), an SO, to gliding *M. gallisepticum* cells. The cell suspension was inserted into the tunnel chamber and observed by phase-contrast microscopy. After 60 s, the buffer was replaced by the buffers containing 0□0.5 mM SL. The gliding cells slowed down after the addition of free SL, and some of them detached from the glass surface (Fig. 3A and B). Most of the cells stopped for gliding kept binding to the end of the membrane protrusion (see Fig. S2 in the supplemental material). The inhibition ratio for binding was calculated from the number of gliding cells at 40 s after the addition of SL to the number before the SL treatments (45). The ratio decreased with SL concentration from 88% ± 11% (n = 14) in 0 mM to 13% ± 11% (n = 21) in 0.5 mM SL (Fig. 3C). The gliding speed at 40 s after the addition of SL also decreased with SL concentration, from 0.17 ± 0.06 μm/s (n = 58) in 0 mM to 0.06 ± 0.04 μm/s (n = 54) in 0.5 mM SL (Fig. 3D). The Hill constant of binding was calculated as previously described (45) to be about 1.55 (see Fig. S3 in the supplemental material), indicating cooperativity in binding between cells and SL.

**Fig. 3.**
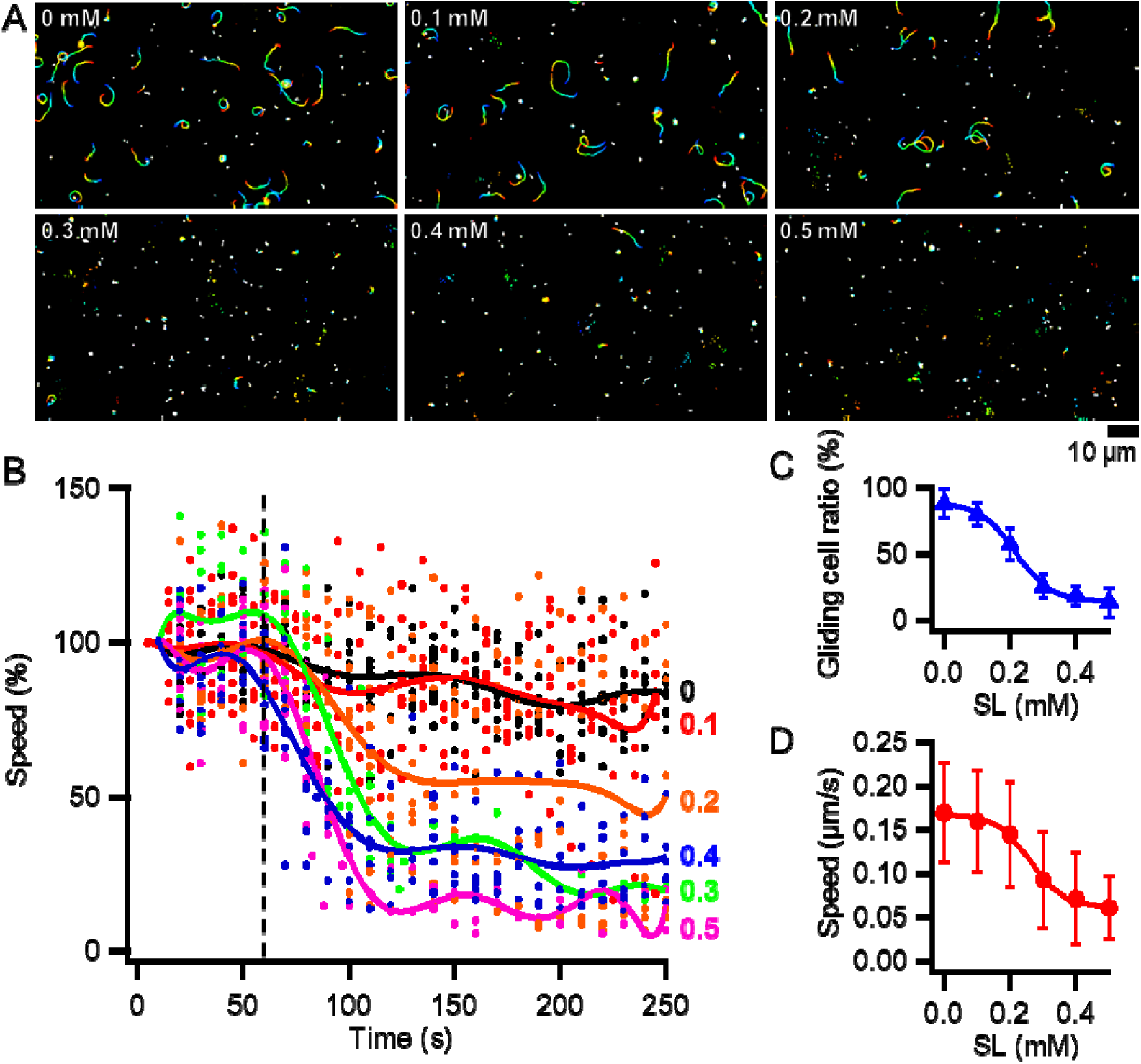
Effects of SL on binding and gliding. (A) Cell trajectories for 60 s under various concentrations of SL. The color changes with time from red to blue. (B) Changes in gliding speeds by the addition of SL. Dotted lines indicate the time point when various concentrations of SL were added. The gliding speeds of individual cells are plotted as dots for every 10 s. (n_cells_ = 9, 9, 10, 6, 6, 9 for 0, 0.1, 0.2, 0.3, 0.4, 0.5 mM SL, respectively). The average gliding speeds are shown with solid lines. The gliding speeds from 0 to 10 s in each condition are normalized as 100%. (C) The gliding cell ratios under each SL concentration are plotted with standard deviations and fitted with a sigmoidal curve. (D) The gliding speeds under each SL concentration are plotted with standard deviations and fitted with a sigmoidal curve.

### Gliding ghost

Previous studies show that the gliding motility of *M. mobile* is driven by ATP hydrolysis based on gliding ghost experiments. In these experiments, *M. mobile* cells were permeabilized with Triton X-100 and stopped for gliding. Gliding was then reactivated by the addition of ATP (32-34). In contrast, the reactivation of permeabilized cells of *M. pneumoniae*-type gliding *Mycoplasmas* has not been successful so far (8). At first, the same method was tried for *M. gallisepticum*, but about half of the cells permeabilized with Triton X-100 detached from the glass surfaces after the addition of ATP solution. Cells were permeabilized with Triton X-100 and ATP to efficiently observe the behaviors of permeabilized cells in the presence of ATP. This strategy was applied to *M. mobile* to confirm whether it was efficient. Cultured *M. mobile* cells were suspended in bufferA (10 mM HEPES pH 7.4, 100 mM NaCl, 2 mM MgCl_2_, 1 mM EGTA, 1 mM DTT, 0.1% MC) containing 20 mM glucose and inserted into the tunnel chamber. Then, the cells were permeabilized with 0.013% Triton X-100 containing 1 mM ATP + 0.01 mM ADP or 1 mM ADP. The cells permeabilized with Triton X-100 containing 1 mM ATP + 0.01 mM ADP glided at a similar speed to the intact cells. The cells permeabilized with Triton X-100 containing 1 mM ADP stopped gliding when they were permeabilized (see Movie S3 and S4 in the supplemental material). These results indicate that this method works efficiently. This method was then applied to *M. gallisepticum*. Cultured *M. gallisepticum* cells were suspended in bufferA containing 10% non-heat-inactivated horse serum, inserted into the tunnel chamber, washed by bufferA containing 20 mM glucose, and observed by phase-contrast microscopy. Under these conditions, the proportion of gliding cells to all cells and the averaged gliding speed were 74% ± 3% (n = 2011) and 0.36 ± 0.06 μm/s (n = 220), respectively (see Fig. S4 in the supplemental material). The buffer was then replaced by 0.007% Triton X-100 containing no ATP, 1 mM ATP + 0.01 mM ADP, or 1 mM ADP in bufferA. The cells became round at 10□30 s from the addition of Triton solutions (Fig. 4A and B) causing the gliding speeds to decrease (Fig. 4C; see Fig. S5 in the supplemental material). The round cells showed three levels of cell-image density shifts; (i) the image density did not decrease, (ii) the image density decreased to be about 75% of the intact cell, (iii) the image density decreased to half of the intact cell (Fig 4D and E; see Fig. S5 in the supplemental material). The cells which decreased to be about 75% of the cell-image density showed slow gliding when they were permeabilized with Triton X-100 containing 1 mM ATP + 0.01 mM ADP (Fig. 4A and C; see Fig. S5 and Movie S5 in the supplemental material). The cells permeabilized with Triton X-100 containing no ATP or 1 mM ADP solution did not show gliding (n_intact_ = 3943 and 2604, respectively) (Fig. 4F). These results indicate that we succeeded in forming gliding ghosts of *M. gallisepticum*. The occurrence ratio of gliding ghosts was calculated to be 0.41% from the numbers of gliding ghosts and intact cells before Triton treatment (n_intact_ = 11528 and n_ghost_ = 47). The gliding speed of ghosts in 1 mM ATP + 0.01 mM ADP was averaged for 150 s at 10 s intervals and found to be 0.014 ± 0.007 μm/s (Fig. 4G). The 62.9% of gliding ghosts continued to glide for 17 min of video recording. In the ATP hydrolysis cycle, the ATP state becomes the ADP or P_i_ state through the ADP + P_i_ state. Vanadate ion (V_i_) with ADP can mimic the ADP + P_i_ state. V_i_ acts as a phosphate analog to form an ADP-V_i_ complex which occupies the catalytic site of ATPase and blocks the hydrolysis cycle (47, 48). The cells were permeabilized with Triton containing 1 mM ATP + 0.5 mM V_i_ to examine whether V_i_ affects the gliding of ghosts, (Fig. 4F). The gliding ghosts in ATP + V_i_ glided with 0.011 ± 0.010 μm/s (n = 9) at an occurrence ratio of ghost to all intact cells 0.37% (n_intact_ = 2439 and n_ghost_ = 9) (Fig. 4H and I), similar to the ghosts constructed by Triton X-100 containing 1 mM ATP + 0.01 mM ADP. However, only 33.3% of ghosts continued to glide for 17 min of video recording under ATP + V_i_ conditions, which is half of the ghosts constructed by Triton X-100 containing 1 mM ATP + 0.01 mM ADP (Fig. 4J), suggesting that V_i_ gradually stopped the gliding of ghosts. These results indicate that the gliding motility of *M. gallisepticum* is driven by ATP hydrolysis.

**Fig. 4.**
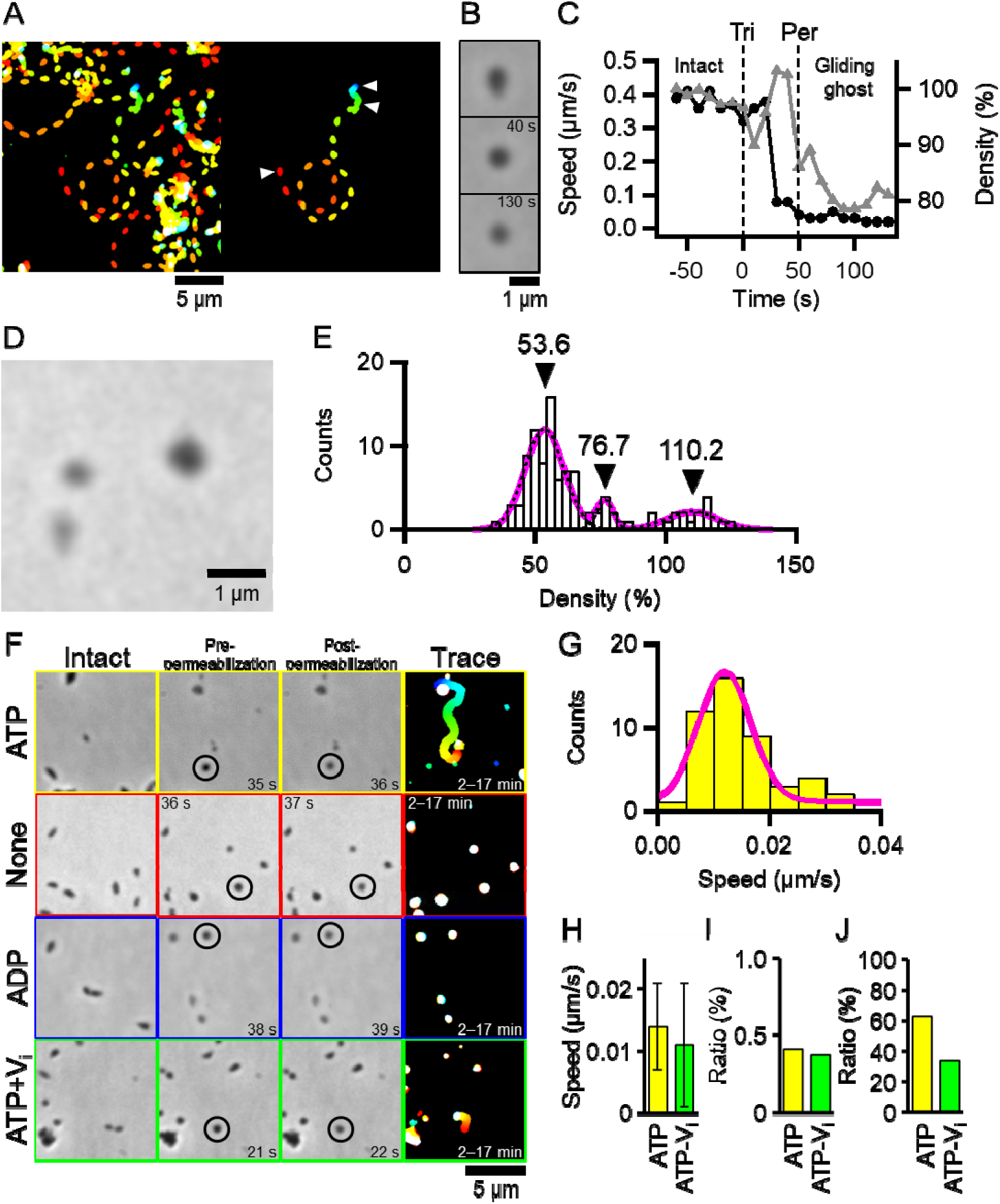
Gliding ghost. (A) Trajectories for 250 s at 5 s intervals of all cells (left) and the cell to be the gliding ghost (right). The color changes with time from red to blue. The 0.007% Triton X-100 containing 1 mM ATP + 0.01 mM ADP solution was added at 0 s. (B) Phase-contrast micrographs of intact (upper), pre-permeabilized (middle) and post-permeabilized cells (lower) marked with white triangles in (A). The time points after the addition of Triton solutions are shown in each panel. (C) Transitions of gliding speeds and cell-image densities. The average gliding speeds and cell-image densities for 10 s at 1-s intervals are plotted as black circles and gray triangles, respectively. The average cell-image densities from −60 to −50 s is normalized as 100%. ‘Tri’ and ‘Per’ indicate the time points when 0.007% Triton X-100 containing 1 mM ATP + 0.01 mM ADP solution was added, and the density shift occurred, respectively. (D) The phase-contrast micrograph of cells treated with Triton solution. Cells showed three levels of cell-image density shifts; the image density did not decrease (right-upper), the image density decreased to be about 75% of the intact cell (left-upper), and the image density decreased to half of the intact cell (left-lower). (E) Distribution of cell-image densities at 90 s after the addition of Triton solution was fitted by the sum of three Gaussian curves. The individual Gaussian curves and the sum of three Gaussian curves are indicated by black broken and magenta solid lines, respectively. The cell-image density before Triton treatment is normalized as 100%. The positions of peak tops are indicated by black triangles. (F) Ghosts constructed by Triton X-100 solutions including nucleotides. Intact, pre-permeabilized, post-permeabilized, and trajectories of ghosts for 15 min are shown from left to right panels. The color changes with time from red to blue. The time points after the addition of Triton solutions are shown in each panel. The cells indicated by black circles decrease for the cell-image densities to be about 75% at post-permeabilized panels. (G) Distribution of gliding speeds of ghosts constructed by 0.007% Triton X-100 containing 1 mM ATP + 0.01 mM ADP solution fitted with a Gaussian curve. (H) The average gliding speeds of ghosts constructed with Triton X-100 solution including ATP or ATP-V_i_ are shown with standard deviations. (I) The occurrence ratios of gliding ghosts to all intact cells are shown. (J) The ratios of ghosts which continued to glide through 17 min of video recording are shown.

### Damage to cell membranes by treatment with Triton X-100

Cells treated with 0.007% Triton X-100 became round and showed three levels of cell-image density shifts (Fig. 4D and E; see Fig. S5 in the supplemental material). Negative-staining electron microscopy was carried out to analyze the morphological changes and cell-image density shifts in detail. Intact cells showed a pear-shape with a membrane protrusion called a ‘bleb’ and ‘infrableb,’ as previously reported (Fig. 5A)(37, 49). Cells treated with Triton X-100 solution containing 1 mM ATP + 0.01 mM ADP showed a round cell shape (Fig. 5B), consistent with our observations by optical microscopy. Some of the cells had large or small holes on the cell membrane (Fig. 5C‒E). These results indicate that the cells treated with Triton X-100 solution have a permeabilized membrane causing the loss of cytoplasm.

**Fig. 5.**
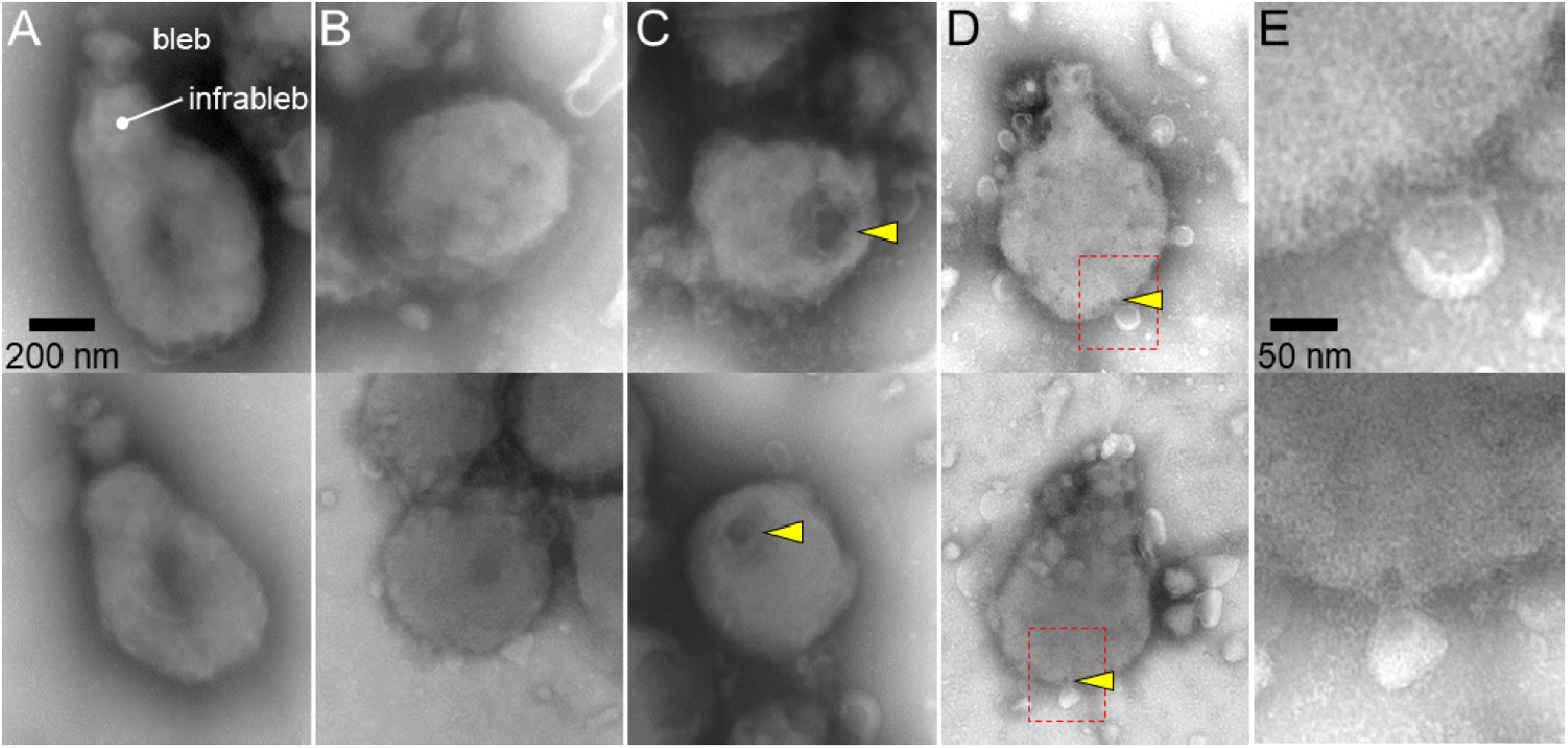
Negative-staining electron micrographs of cell permeabilization. (A) Intact cells. The cells featured by the bleb and infrableb are shown. (B–E) Cells treated with 0.007% Triton X-100 containing 1 mM ATP + 0.01 mM ADP. Some of the permeabilized cells have large (C) or small (D) holes marked by yellow triangles. (E) Magnified images of red boxed areas in (D). A–D show the same magnification. Two examples of images are shown for each category.

## DISCUSSION

### Gliding behaviors

In the present study, we observed and analyzed the gliding behaviors of *M. gallisepticum* by using optical microscopes. The gliding speed of *M. gallisepticum* measured in the present study was about 0.27 μm/s (Fig. 1C), which is comparable to the reported gliding speeds of *M. pneumoniae* and *M. genitalium*, about 0.64 μm/s and 0.15 μm/s, respectively (39). A previous study shows that *M. genitalium* has curved attachment organelles and a circular trajectory of gliding, but the deletion mutant of MG_217 protein shows straight attachment organelles and a straight trajectory, suggesting that the gliding direction is determined by the alignment of the attachment organelle (23). *M. gallisepticum* has non-bended attachment organelles (Fig. 5A) (37), and the average gliding direction was 0.6 ± 44.6 degrees/5 s (Fig. 1D), which is consistent with previous observations (23, 41). However, cells sometimes glided to the left or the right (Fig. 1B). In these cases, the cells bind to glass surfaces at the end of the membrane protrusion so that the thermal fluctuations and drag of the cell body likely changes the gliding direction to left or right.

*M. gallisepticum* did not roll around the cell axis during gliding (Fig. 1E; see Fig. S1 in the supplemental material). Previous studies show that *M. mobile* does not roll even with adhesin complexes working as a ‘leg’ around the membrane protrusion (41, 50, 51). *M. gallisepticum* and *M. mobile* glide on the ciliated epithelial cells of birds and the gills of fresh-water fish in nature, respectively. The distribution of adhesin complexes may be an advantage for binding to host surfaces because these tissue surfaces are three-dimensionally aligned.

### Different inhibitory effects of MC and Ficoll on gliding

In this study, it was found that MC and Ficoll have different inhibitory effects on *M. gallisepticum* gliding (Fig. 2). Previous studies show that a periplasmic flagellated bacterium whose flagella are located between the inner and outer membranes, *Brachyspira pilosicoli*, exhibits swimming at a constant speed in various concentrations of polyvinylpyrrolidone. These results show similar characteristics to MC, but the swimming speed decreased with Ficoll concentrations (43). External flagellated bacteria, *Pseudomonas viscosa*, *Bacillus brevis,* and *Escherichia coli*, increased swimming speeds in 1.33 mPa s of MC (52). Magariyama and Kudo suggested in 2002 that a gel-like three-dimensional network formed by MC makes a virtual tube around the cell body, and the body moves easily in the tube (42). This virtual tube may support the gliding motility of *M. gallisepticum*, enabling the cells to glide even in viscous environments constantly.

### Cooperativity in binding

Binding and gliding speed sigmoidally decreased by free SL (Fig. 3C and D). Previous studies show that one adhesin complex of *M. pneumoniae* is composed of two heterodimers, one of each is assembled by one P1 adhesin molecule and one P90 molecule (28). The adhesin complex in *M. genitalium* is also composed of a dimer of heterodimers constructed by P110 and P140, the homologs of P1 adhesin and P90, respectively (53). P110 has a binding site for SOs, so one adhesin complex binds two SOs (31). The adhesin complex of *M. gallisepticum* is assumed to have two binding sites for SOs because it is composed of CrmA and GapA, the homologs of P110 and P140, respectively. This assumption is consistent with 1.55 of the Hill constant. The Hill constant of binding between *M. pneumoniae* cells and sialic compounds ranges from 1.5 to 2.5 (45), comparable to 1.55 of the Hill constant for *M. gallisepticum* (see Fig. S3 in the supplemental material).

In 0.5 mM SL conditions, cells rotating around the end of the membrane protrusion were observed (see Fig. S2 in the supplemental material). A previous study shows that the ghosts of *M. mobile* exhibit directed rotational motility around the membrane protrusion driven by the linear motion of the legs (34). In contrast, rotary motion in *M. gallisepticum* seems to be driven by thermal fluctuation because it has no regularity of rotational direction (see Fig. S2 in the supplemental material), suggesting the possibility that adhesin complexes exist with high density at the end of the membrane protrusion.

### Energy source

In the present study, we succeeded in forming the gliding ghosts of *M. gallisepticum* and clarified that the direct energy source of gliding is ATP (Fig. 4). In this method, we added 0.01 mM ADP to 0.007% Triton X-100 and 1 mM ATP solution because cells permeabilized with 0.007% Triton X-100 and 1 mM ATP solution easily detach from glass surfaces and ADP reduces these detachments.

In a previous study, the gliding ghosts of *M. mobile* glided at similar speeds to intact cells, and 85% of ghosts showed gliding (32). However, the gliding speed of *M. gallisepticum* ghosts was 4% of that of the intact cells, and 0.4% of the intact cells became gliding ghosts (Fig. 4I and J). Kawamoto *et al*. showed in 2016 that the translucent area surrounding the internal core in *M. pneumoniae* might be occupied by less-di□usive materials and play a role in transmitting the movements of the paired plates originating in the bowl complex to the adhesin complexes (7). The low occurrence ratio of gliding ghosts in *M. gallisepticum* may be caused by the permeabilization of cells resulting in the elution of less-di□usive materials. The cells treated with Triton X-100 show three levels of cell-image density shifts (Fig. 4D and E). The cells which decreased in the cell-image density to be 75% of the intact cells probably have permeabilized cell membranes which retain most of the less-di□usive materials.

In the gliding motility of *M. mobile*, complexes of MMOBs 1660 and 1670 in internal jellyfish-like structures have been proposed to hydrolyze ATP molecules as a motor (5, 54). Therefore, which proteins work as a motor in the *M. pneumoniae*-type gliding system need to be identified. Fifteen component proteins of the attachment organelles in *M. pneumoniae* have been identified until now (8). One of these proteins has been annotated as Lon, an ATP-dependent protease (8). This protein possibly works as a motor for gliding.

Generally, respirable bacteria generate a proton gradient across the cell membrane in the respiratory process. The proton gradient causes proton motive force which drives F-type ATP synthase and bacterial flagella (55, 56). However, *Mycoplasmas* have no genes for electron transport and synthesize ATP molecules by glycolysis (57). The membrane potential of *M. gallisepticum* was −48 mV, much smaller than that of typical bacteria, which is −150 mV (58-61). Therefore, ATP is more convenient for *Mycoplasmas* for the energy source of gliding motilities rather than proton motive force.

## MATERIALS AND METHODS

### Cultivation

The *M. gallisepticum* S6 strain was grown in Aluotto medium at 37□, as previously described (37).

### Observations of gliding behaviors

The cells were cultured to reach an optical density at 595 nm of around 0.1. The cultured cells were collected by centrifugation at 12,000 × *g* for 10 min at room temperature (RT) and suspended in PBS consisting of 75 mM sodium phosphate (pH 7.3) and 68 mM NaCl. The cell suspension was centrifuged at 12,000 × *g* for 5 min at RT, suspended in PBS containing 10% non-heat-inactivated horse serum (Gibco; Thermo Fisher Scientific, Waltham, MA), poured through a 0.45-μm pore size filter and incubated for 15 min at RT. Then, the cell suspension was poured twice more and inserted into a tunnel chamber which was assembled by taping coverslips cleaned with saturated ethanolic KOH and precoated with 100% non-heat-inactivated horse serum for 60 min and 10 mg/ml bovine serum albumin (Sigma-Aldrich, St. Louis, MO) in PBS for 60 min at RT. The tunnel chamber was washed with PBS containing 20 mM glucose, incubated at 37□ on an inverted microscope (IX83; Olympus, Tokyo, Japan) equipped with a thermo plate (MATS-OTOR-MV; Tokai Hit, Shizuoka, Japan) and a lens heater (MATS-LH; Tokai Hit), observed by phase-contrast microscopy at 37□ and recorded with a charge-coupled device (CCD) camera (LRH2500XE-1; DigiMo, Tokyo, Japan). Video data were analyzed by ImageJ 1.43u (http://rsb.info.nih.gov/ij/), as previously described (37, 41).

To investigate the effect of viscosity on gliding, the cultured cells were washed with PBS containing 10% non-heat-inactivated horse serum and 0.10%, 0.25%, and 0.50% MC (Methyl Cellulose #400; nacalai tesque, Kyoto, Japan) or 5%, 10%, and 15% Ficoll (M.W. 400,000; nacalai tesque, Kyoto, Japan), poured and incubated. Then, the cell suspension was poured and inserted into a cleaned and precoated tunnel chamber. The tunnel chamber was washed with various concentrations of viscous buffer containing 20 mM glucose, observed by phase-contrast microscopy at 37□ and recorded. The viscosities were measured using dynamic viscoelasticity measuring apparatus (Rheosol-G5000; UBM, Kyoto, Japan) at 37□ as follows: 0.66 mPa s for PBS/G, 2.5, 5.5, 7.3 mPa s for 0.10%, 0.25%, 0.50% MC, 2.3, 3.0, 4.6 mPa s for 5%, 10%, 15% Ficoll.

Cells on the tunnel chamber were treated with various concentrations of 3′-sialyllactose sodium salt (Nagara Science Co., Ltd, Tokyo, Japan) in PBS/G, observed by phase-contrast microscopy at 37□ and recorded to examine the binding features. The cultured cells were collected, suspended in PBS containing 10 mM Sulfo-NHSLC-LC-Biotin (Thermo Fisher Scientific) and incubated for 15 min at RT to observe cell rolling. The cell suspension was centrifuged, suspended in PBS containing 10% non-heat-inactivated horse serum, poured, and incubated. Then, the cell suspension was poured and inserted into a cleaned and precoated tunnel chamber. The tunnel chamber was washed by PBS/G containing streptavidin conjugated 40 nm colloidal gold (Cytodiagnostics, Ontario, Canada), observed by dark-field microscopy using an upright microscope (BX50; Olympus) at 37□ and recorded by a CCD camera (WAT-120N; Watec Co. Ltd., Yamagata, Japan). Video data were analyzed by ImageJ 1.43u and IGOR Pro 6.33J (WaveMetrics, Portland, OR).

### Gliding ghost

The cultured cells were collected and suspended in HEPES buffer (10 mM HEPES pH 7.4, 100 mM NaCl). The cell suspension was centrifuged, suspended in bufferA, poured, and incubated for 15 min at RT. Then, the cell suspension was poured and inserted into a cleaned and precoated tunnel chamber. The tunnel chamber was washed with bufferA containing 20 mM glucose, observed by phase-contrast microscopy at 37□, and recorded. After 70 s, the buffer was replaced with 0.007% Triton X-100 (MP Biomedicals, Santa Ana, CA) containing 1 mM ATP and 0.01 mM ADP or 0 mM ATP or 1 mM ADP or 1 mM ATP and 0.5 mM Na_3_VO_4_. Video data were analyzed by ImageJ 1.43u.

### Negative-staining electron microscopy

The cultured cells were collected, suspended in PBS containing 10% non-heat-inactivated horse serum, poured through a 0.45-μm pore size filter and incubated for 15 min at RT. Then, the cell suspension was placed on a carbon-coated grid and incubated for 10 min at RT. The grid was treated with 0.007% Triton X-100 containing 1 mM ATP and 0.01 mM ADP to permeabilize cells, incubated for 1 min at RT and fixed with 1% glutaraldehyde in bufferA for 1 min at RT. The fixed cells were washed with water, stained with 0.5% ammonium molybdate, and observed using a transmission electron microscope (JEM-1010; JEOL, Tokyo, Japan) at 80 kV equipped with a CCD camera (FastScan-F214 (T); TVIPS, Gauting, Germany).

## ACKNOWLEDGMENTS

We thank Yuhei O Tahara at Osaka City University for technical assistance with negative-staining electron microscopy. This work was supported by a Grant-in-Aid for Scientific Research in the innovative area of ‘’Harmonized Supramolecular Motility Machinery and Its Diversity’ (Ministry of Education, Culture, Sports, Science, and Technology KAKENHI; grant number 24117002) and by a Grants-in-Aid for Scientific Research (A) (Ministry of Education, Culture, Sports, Science, and Technology KAKENHI; grant number 17H01544) to M. Miyata. M. Mizutani is the recipient of a Research Fellowship of the Japan Society for the Promotion of Science (18J15362).

---

**Fig. S1.**
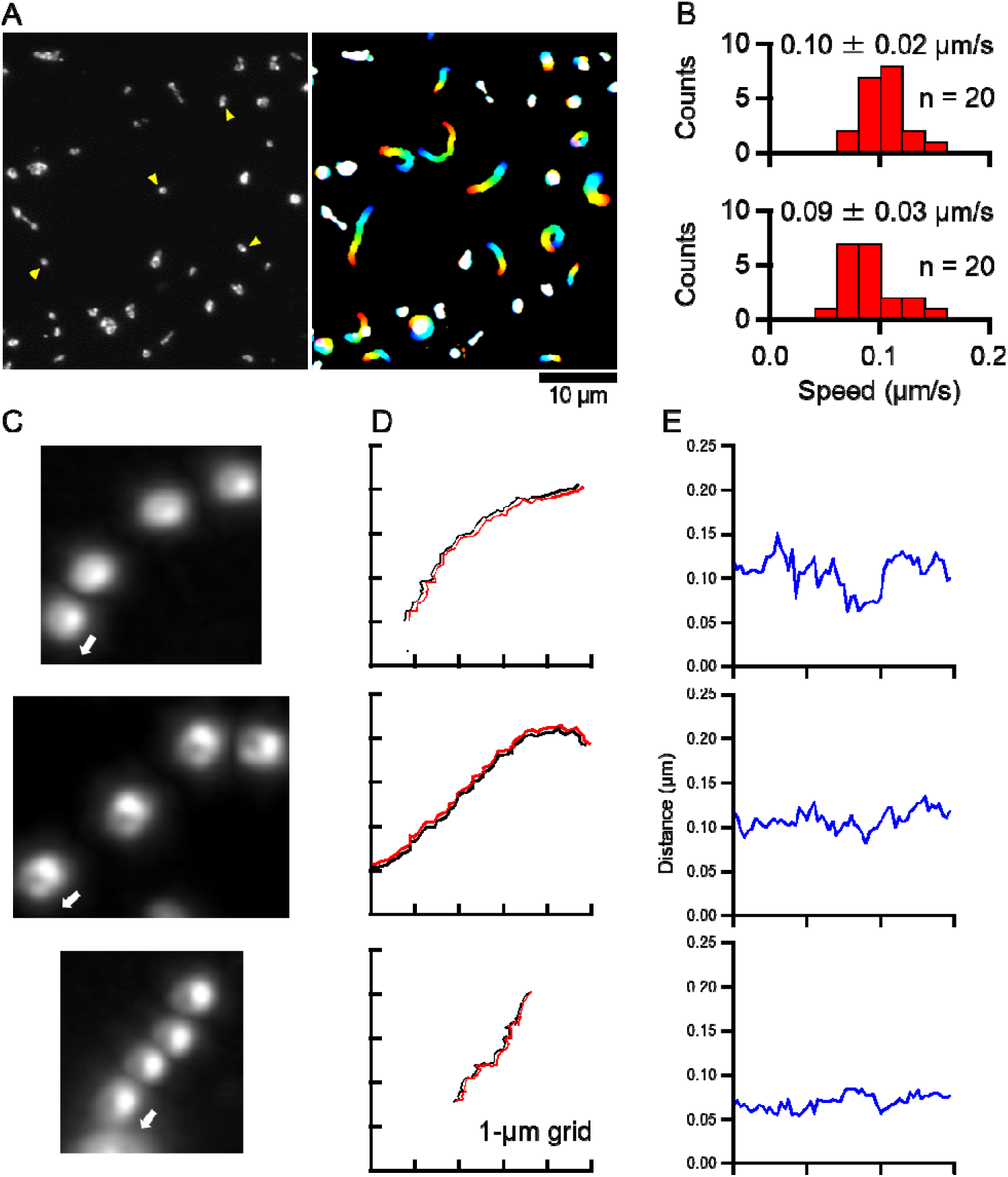
Gliding motility of cells attached with colloidal gold. (A) The field image of dark-field microscopy. The yellow triangles indicate the examples of colloidal gold attached to cells. (left). The cell trajectories for 60 s, changing color with time from red to blue (right). (B) The distributions of gliding speeds with (upper) and without colloidal gold (lower). The averaged gliding speeds with standard deviations are shown in each panel. (C). Four consecutive dark-field micrographs of a cell attached with a colloidal gold at 20 s intervals are shown in the same field. The cell glided in the direction indicated by a white arrow. These micrographs are shown in scale with (D). (D) The trajectories of the mass centers of cells and colloidal gold are indicated by black and red lines, respectively. (E) Distances between the mass centers of cells and colloidal gold are indicated by blue lines.

**Fig. S2.**
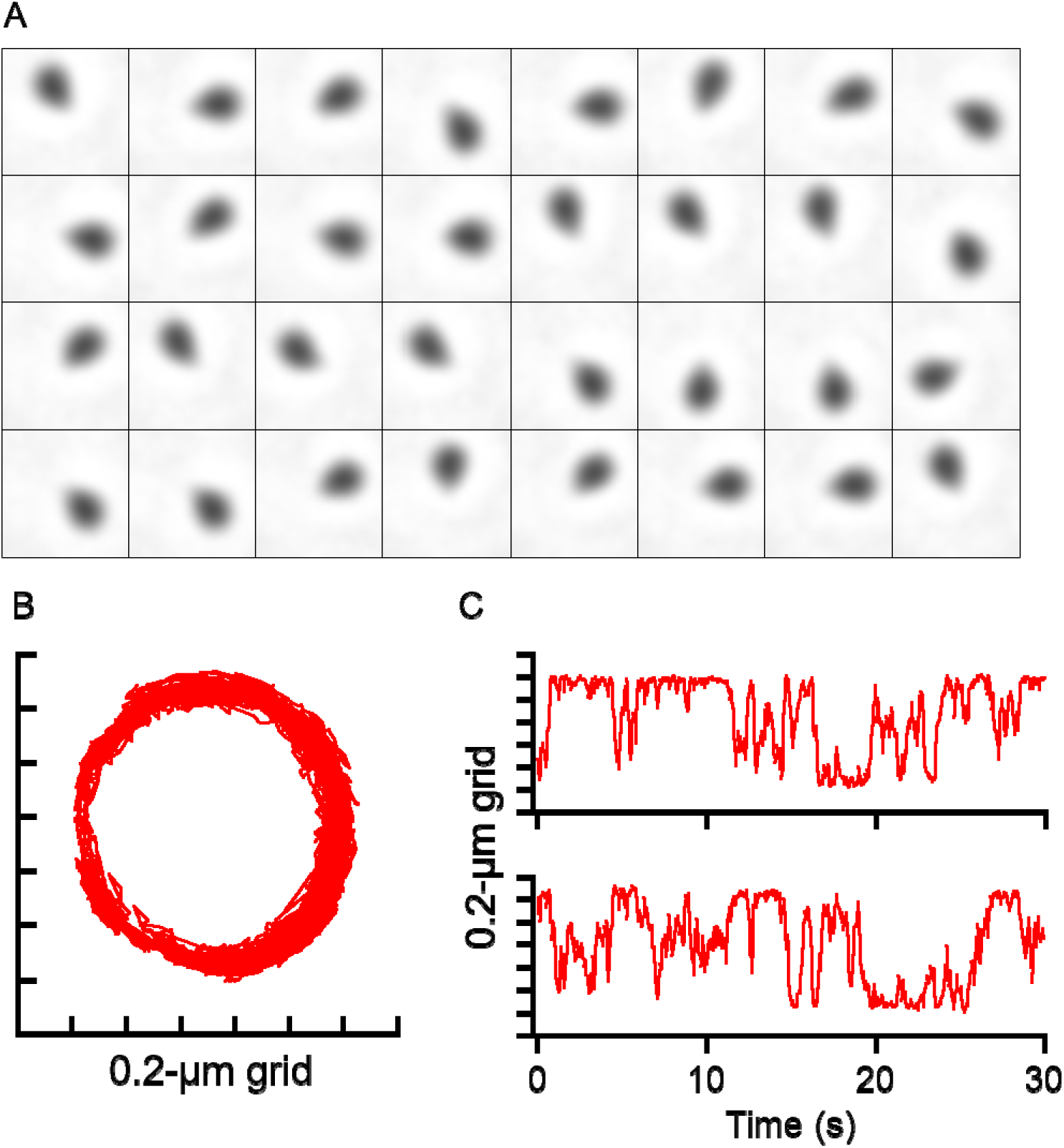
Cells rotating around the end of the membrane protrusion under 0.5 mM SL. (A) Sequential phase-contrast micrographs of a cell at 1 s intervals. (B) The mass center of the cell was traced at 5 ms intervals for 30 s. (C) Time courses against the *X*-axis (upper) and *Y*-axis (lower) of panel B.

**Fig. S3.**
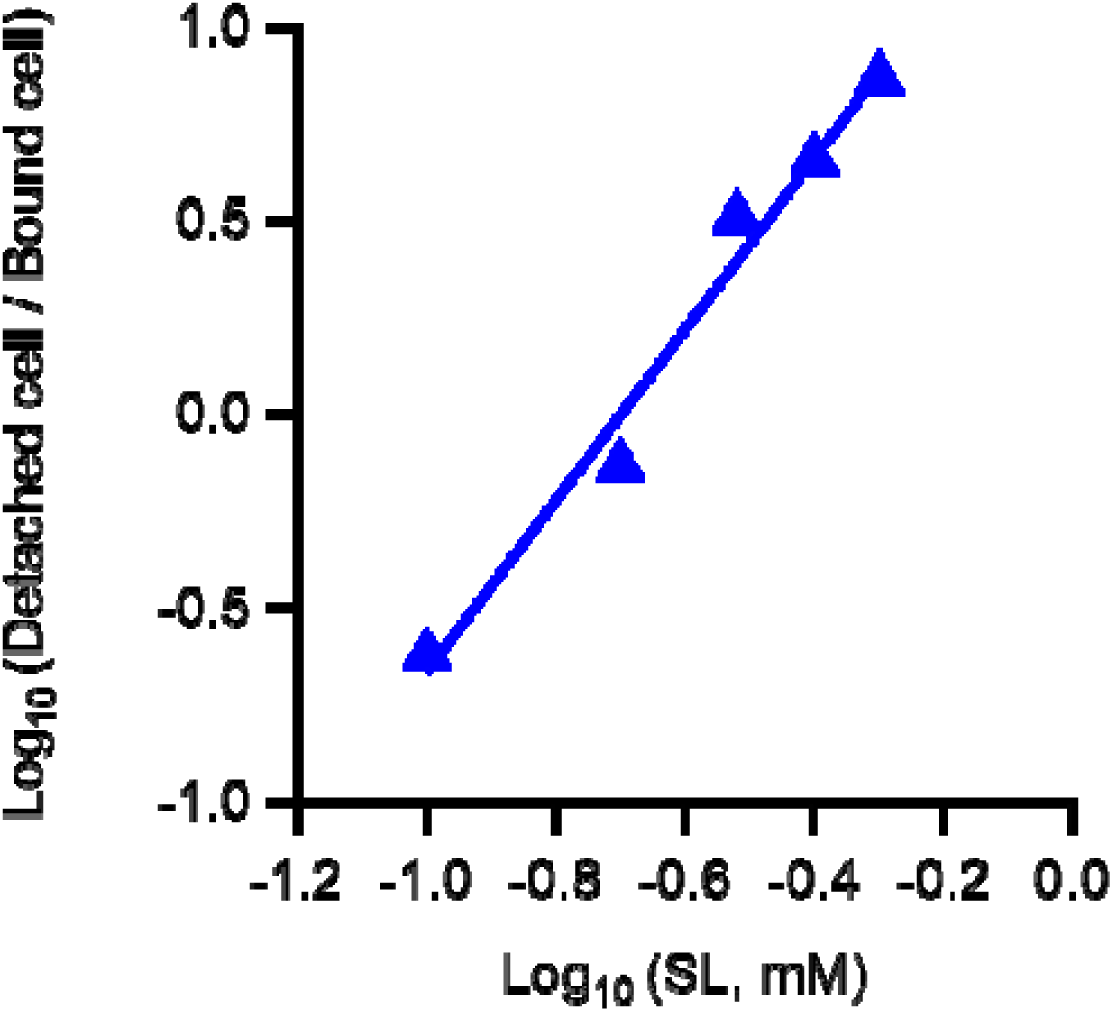
Hill plot analysis of data in Fig. 3C.

**Fig. S4.**
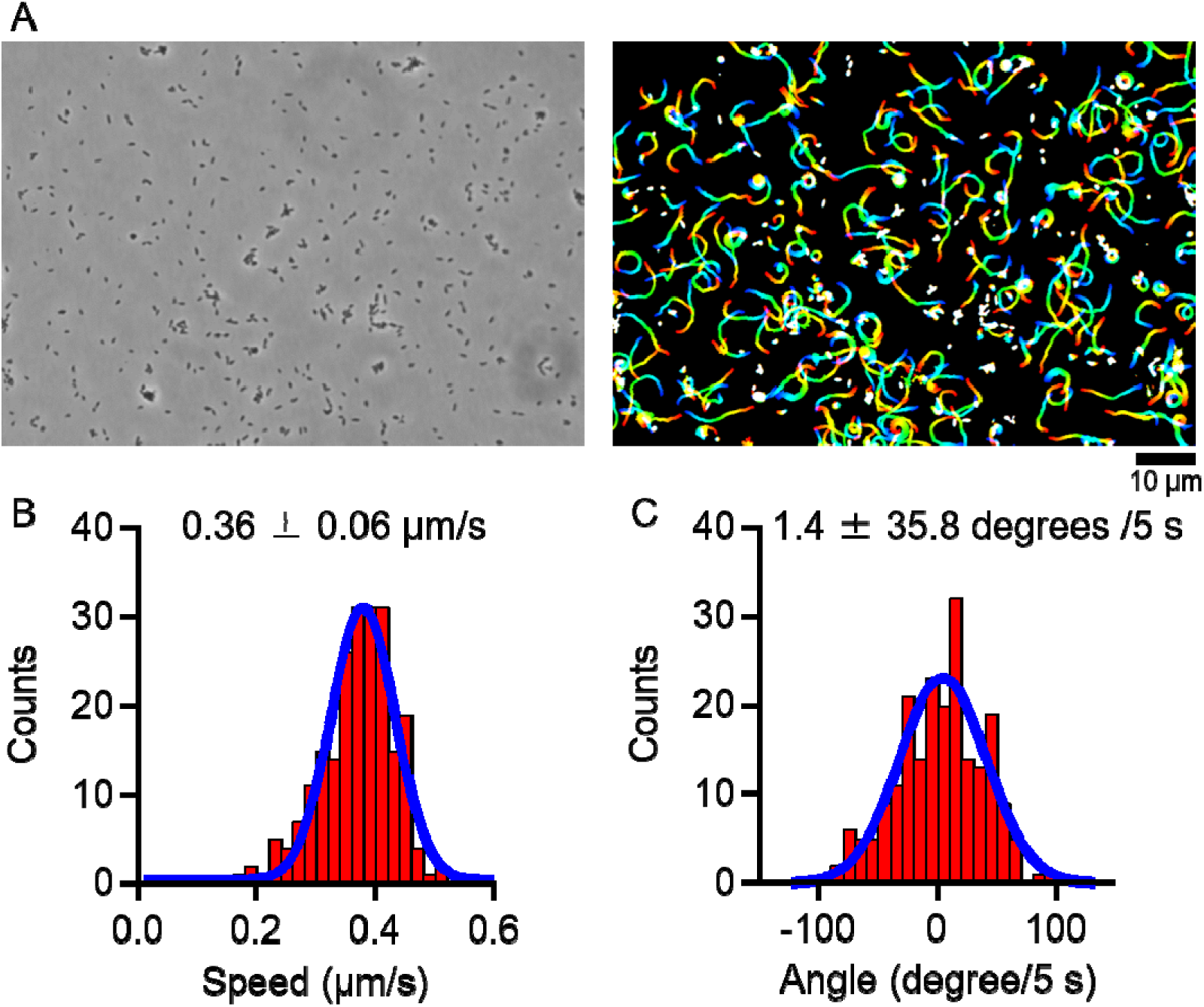
Gliding motility under bufferA. (A) Field image of phase-contrast micrograph (left) and cell trajectories for 30 s, changing color with time from red to blue (right). (B) Distribution of gliding speeds averaged for 60 s under bufferA was fitted with a Gaussian curve. (C) Distribution of gliding directions under bufferA was fitted with a Gaussian curve.

**Fig. S5.**
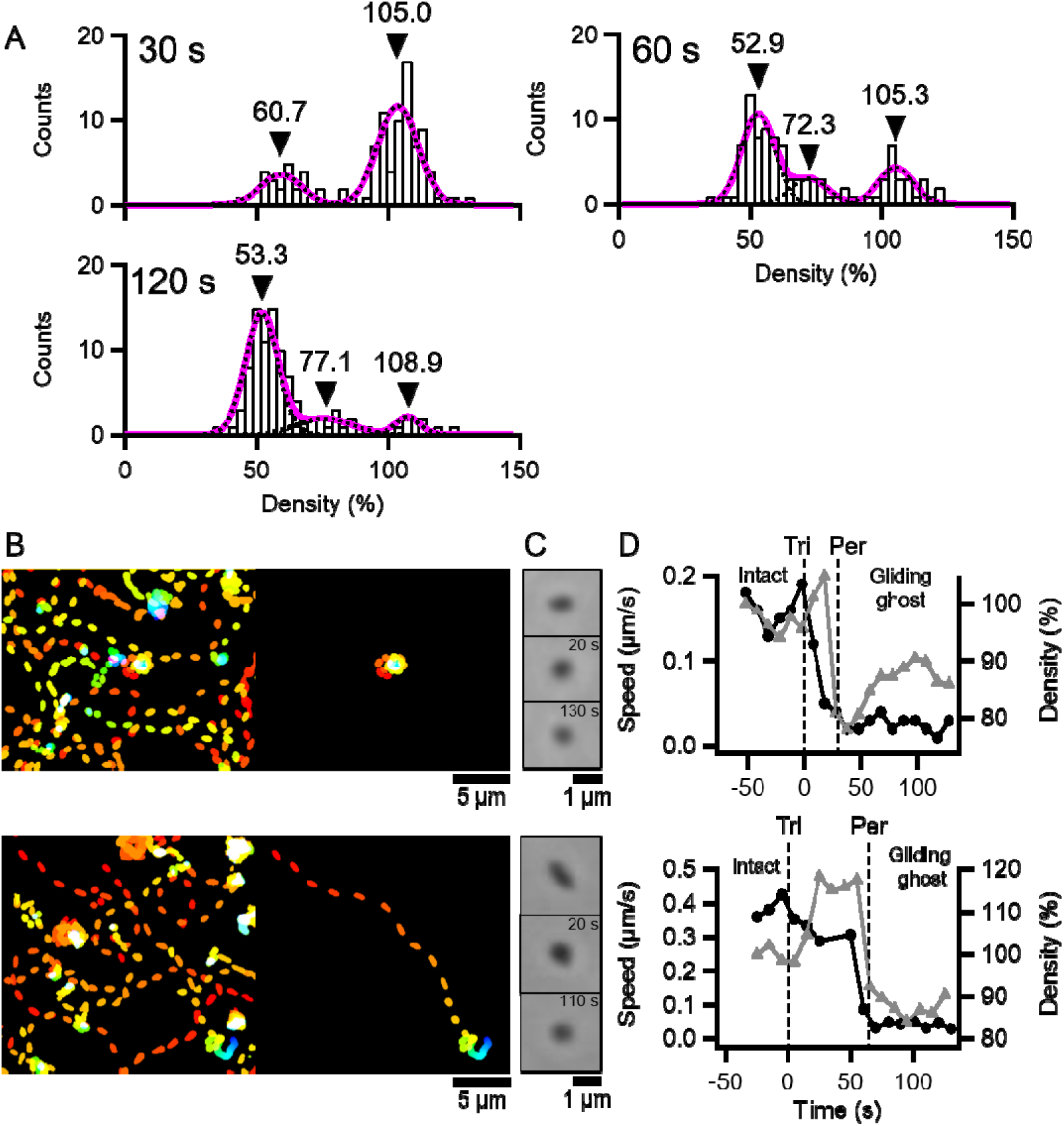
Transition of cell-image density in gliding ghost experiments. (A) Distributions of cell-image densities at 30, 60, and 120 s after the addition of Triton solution was fitted by the sum of two or three Gaussian curves. The individual Gaussian curves and the sum of the Gaussian curves are indicated by black broken and magenta solid lines, respectively. The cell-image density before Triton treatment is normalized as 100%. The positions of peak tops are indicated by black triangles. (B) Trajectories at 5 s intervals of all cells (left) and the cell to be the gliding ghost (right). The color changes with time from red to blue. The 0.007% Triton X-100 containing 1 mM ATP + 0.01 mM ADP solution was added at 0 s. (C) Phase-contrast micrographs of intact (upper), pre-permeabilized (middle) and post-permeabilized cells (lower). The time points after the addition of Triton solutions are shown in each panel. (D) Transitions of gliding speeds and cell-image densities. The average gliding speeds and cell-image densities for 10 s at 1 s intervals are plotted as black circles and gray triangles, respectively. The average cell-image densities of the first 10 s are normalized as 100%. ‘Tri’ and ‘Per’ indicate that the time points when 0.007% Triton X-100 containing 1 mM ATP + 0.01 mM ADP solution was added, and the density shift occurred, respectively. Two examples are shown for each category.

Movie S1

Gliding *M. gallisepticum* under normal conditions. The movie is 5× speed.

Movie S2

Gliding *M. gallisepticum* attached with colloidal gold. The movie is 5× speed.

Movie S3

Gliding ghost of *M. mobile* under ATP. The movie is 2× speed.

Movie S4

Gliding ghost of *M. mobile* under ADP. The movie is 2× speed.

Movie S5

Gliding ghost of *M. gallisepticum* under ATP. The movie is 30× speed.

